# The Y951N patient mutation inactivates the intramolecular switch in human mitochondrial DNA POLγ

**DOI:** 10.1101/2024.08.28.610080

**Authors:** Josefin M. E. Forslund, Tran V.H. Nguyen, Vimal Parkash, Andreas Berner, Steffi Goffart, Jaakko L.O. Pohjoismäki, Paulina Wanrooij, Erik Johansson, Sjoerd Wanrooij

## Abstract

Mitochondrial DNA (mtDNA) stability, essential for cellular energy production, relies on DNA polymerase gamma (POLγ). Here, we show that the POLγ Y951N disease causing mutation induces replication stalling and severe mtDNA depletion. However, unlike other POLγ disease causing mutations, Y951N does not directly impair exonuclease activity and only mildly affects polymerase activity. Instead, we found that Y951N compromises the enzyme’s ability to efficiently toggle between DNA synthesis and degradation, and is thus the first patient-derived mutation with impaired polymerase-exonuclease switching. These findings provide new insights into the intramolecular switch when POLγ proofreads the newly-synthesized DNA strand, and reveal a new mechanism for causing mitochondrial DNA instability.

**Significance Statement:** DNA polymerase gamma (POLγ) is essential for copying mitochondrial DNA (mtDNA), which is crucial for our energy production. POLγ must accurately switch between making new DNA (polymerase activity) and correcting errors (exonuclease activity). While it is known that mutations in POLγ can cause mitochondrial diseases by directly impairing these enzymatic functions, this study reveals a new mechanism. The Y951N mutation disrupts POLγ’s ability to switch between these activities, leading to severe blockages in DNA replication and a loss of mtDNA in human cells, even without significant direct impairment of polymerase or exonuclease activities. These findings provide new insights into the origins of mitochondrial diseases.

## Introduction

Mitochondria are intracellular organelles essential for energy production. In contrast to other subcellular organelles, the mitochondria contain their own genome called the mitochondrial DNA (mtDNA). Defects in mtDNA maintenance lead to disturbed energy production and can cause a broad range of inherited diseases, neurodegeneration and aging (1).

Human mtDNA is replicated by a unique set of proteins that are distinct from the nuclear DNA replication machinery. DNA polymerase gamma (POLγ) is the replicative DNA polymerase in the mitochondria and is organized as a heterotrimer, consisting of one catalytic subunit (POLγA) and two processivity subunits (POLγB_2_) (2). During DNA replication, POLγA carries out DNA synthesis in the 5’-to-3’ direction and removes its own replication errors with a 3’-to-5’ exonuclease activity. The polymerase and exonuclease sites are separated by 32 Å (3), necessitating that the proofreading activity both recognize a wrongly incorporated nucleotide (mismatch) and transfers the 3’-end from the polymerase site to the exonuclease site. After removing the mismatch in the exonuclease site, the 3’-end of the nascent strand is returned to the polymerase site (4).

Mutations in the *POLG* gene, encoding for the catalytic subunit, are associated with several mitochondrial disorders such as mitochondrial DNA depletion syndrome, progressive external ophthalmoplegia (PEO) and Alpers syndrome (5). Mutations in the polymerase domain of POLγA primarily compromise DNA synthesis, leading to defective mtDNA replication (2, 6, 7), whereas mutations in the exonuclease domain predominantly impair the proofreading function, resulting in an increased accumulation of mtDNA mutations and associated mitochondrial pathologies (8, 9).

In fact, reducing either of the two activities in POLγ will have detrimental effects by affecting the balance between its polymerase and exonuclease function and thus influence the fidelity of DNA replication (10). A shift in this equilibrium, favoring one activity over the other, can lead to a polymerase with either a mutator or an antimutator phenotype (10-12). DNA polymerases continuously shuffle the 3’-end of the nascent DNA-strand between their exonuclease and polymerase active sites. This transition can occur through an intermolecular mechanism, where the polymerase dissociates from the DNA and subsequently reassociates with the DNA bound to a different domain. However, this intermolecular switching process tends to be slower (13) compared to a second mechanism known as the intramolecular switch. During intramolecular switching, the polymerase remains associated with the DNA throughout the entire process, while melting the duplex DNA at the 3’-terminus and only transferring the nascent strand between the polymerase and exonuclease active sites. The mechanistic details of intramolecular switching vary substantially both between and within different DNA polymerase families, lacking a uniform mechanism (14-22).

In recent years, substantial advancements have been made in understanding the intramolecular switching mechanism of human POLγ. Both computational analyses and investigation of the POLγ 3D structure have revealed a dynamic bidirectional communication between the polymerase and exonuclease sites during replication (23-25). This communication is not mediated by direct site-to-site contact, but rather via a sequence of subtle conformational changes within each domain during the switching process. These structural changes alter the polymerase architecture, allowing an active transition between its two activities (23-25). While many patient mutations in POLγA have been found to impair either the polymerase or the exonuclease activity, it remains unexplored how different patient mutations affect the intramolecular switch between the two active sites in POLγ.

In this study, we show that the dominant patient mutation Y951N, located in the polymerase domain of POLγA (26, 27), impairs the intramolecular switch of human mtDNA POL*γ in vitro* and leads to severe mtDNA replication stalling in cells. Our findings indicate that certain patient mutations may not only affect specific catalytic activities directly but also influence the switch between the polymerase site and exonuclease site. The identification of POLγA residues involved in the intramolecular switching mechanism will provide valuable insights into how other patient mutations may impact the maintenance of the mitochondrial genome.

## Results

### POLγ A Y951N expression causes mtDNA replication stalling and depletion in human cells

The POLγA Y951N variant was found in heterozygous state in a patient suffering from bilateral congenital cataracts, ovarian dysgenesis, marked muscle weakness and atrophy (26, 27). The Y951 residue is situated in Motif B of the polymerase domain of POLγA (from now on referred to as POLγ) and is conserved in other Family A DNA polymerases (Fig. 1A-B). This specific tyrosine has previously been shown to play an important role in the polymerase active site, particularly in the interaction with bound dNTPs (3, 28, 29). However, the mechanistic impact of the Y951N amino acid substitution in POLγ is unexplored.

**Figure 1.**
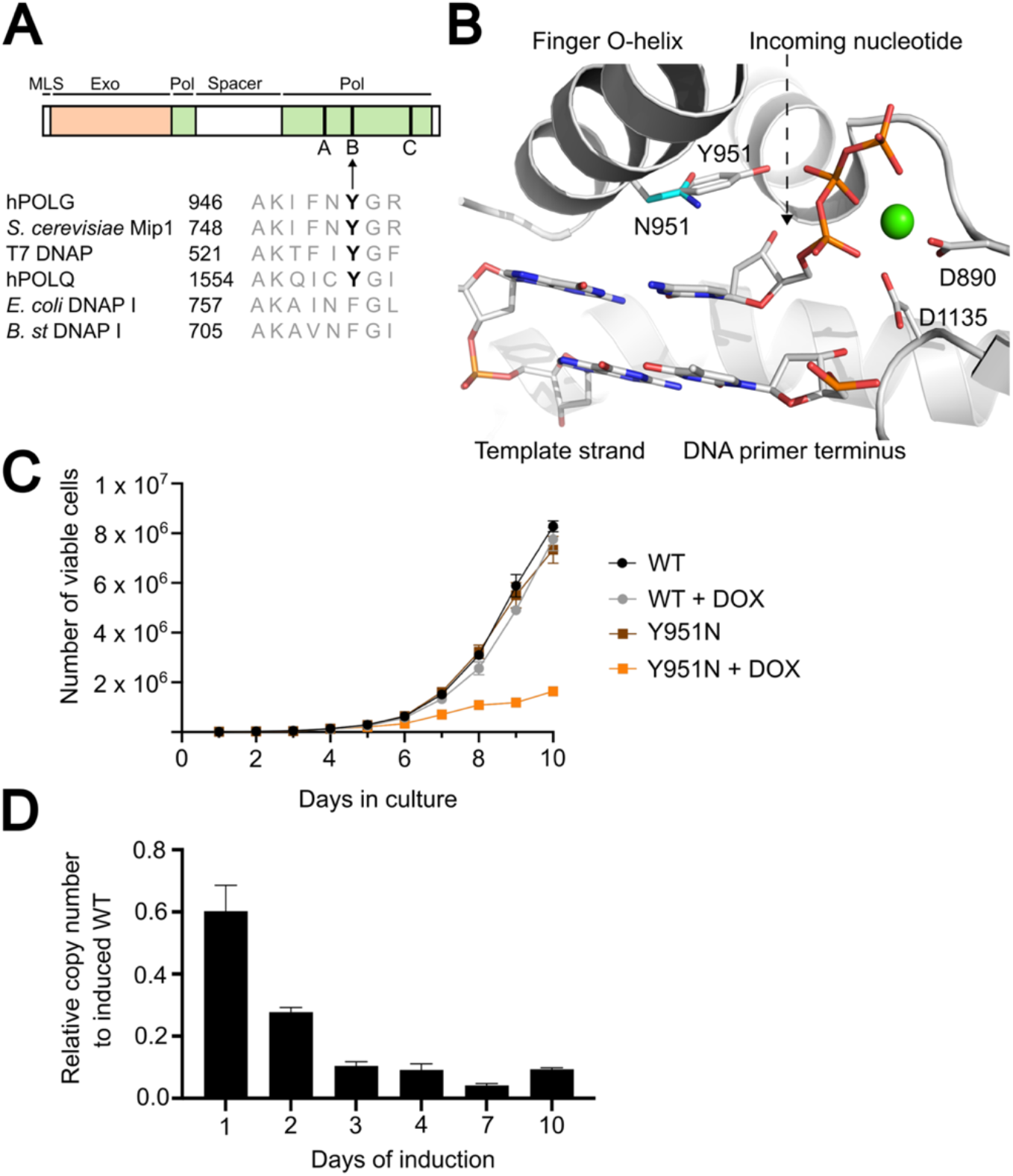
POLγ Y951N patient mutation leads to slow cell growth and decreased mtDNA copy number *in vivo*. **(A)**A linear presentation of POLγ A protein highlighting the tyrosine 951 (Y951) residue located in the B motif of the polymerase domain. Alignment of human POLγ (hPOLG) with other Family A polymerases, including mitochondrial DNA polymerase in *Saccharomyces cerevisiae* (Mip1), T7 DNA polymerase (T7 DNAP), human POLθ (hPOLQ), *Escherichia coli* DNA polymerase I (*E. coli* DNAP I) and *Bacillus stearothermophilus* DNA polymerase I (*B. st* DNAP I), shows that the tyrosine is conserved across several species. MLS; mitochondrial localization signal, Exo; exonuclease domain, Pol; polymerase domain. **(B)**The structure of human DNA POLγ replication complex while extending a G-T mismatch (PDB 8d37; Park et al., 2023). The structure shows the incoming nucleotide in the active site pocket, which is located adjacent to the 3’-primer terminus end. The model illustrates Y951 positioned near the triphosphate group of the incoming nucleotide. Modelling an asparagine at 951 (Y951N) results in a wider gap between N951 and the triphosphate of the incoming nucleotide. The Ca^2+^ (green sphere) at the B-site coordinates the triphosphate and the catalytic aspartates. **(C)**Growth curve of WT POLγ A or Y951N Flp-In T-REx cells in absence (black and brown lines) or presence (grey or orange lines) of 100 ng/ml doxycycline as an expression inducer. Initially, 5 × 10^3^ cells were seeded onto 6-well plates and cultivated for the indicated period. Cells were stained with Trypan blue dye, and viable cells were counted using an automated cell counter. Graph presents total viable cell count as a function of time (days in culture), with error bars represent mean -/+ SD from three independent experiments. **(D)** The relative mtDNA copy number in Flp-In T-REx cells expressing POLγ A Y951N, quantified through multiplex qPCR of totDNA. For quantification, the ratio between mtDNA D-loop region and nuclear DNA B2M region was determined. The relative mtDNA copy number of POLγ A Y951N induced cells is normalized to that of WT POLγ A induced cells. Error bars represent mean -/+ SD of four individual experiments.

To investigate the impact of the Y951N mutation on mtDNA replication within human cells, we generated stable inducible Flp-In T-REx 293 cell lines that express either the wildtype (WT) POLγ-flag protein or its Y951N variant upon addition of doxycycline. Expression of the transgene was verified by Western blot analysis using an anti-flag antibody (Fig. S1A). After a ten-day period of expression, we observed a pronounced impact on the growth of the cells expressing the Y951N mutant compared to that of the WT POLγ-expressing cells (Fig. 1C and Fig. S1B).

Moreover, multiplex qPCR analysis revealed a notable progressive decrease in mtDNA copy number when overexpressing the Y951N variant in cells (Fig. 1D). Additionally, 2D gel analysis confirmed that the observed mtDNA depletion coincided with marked replication stalling in the mutant induced cells, evident from the accumulation of mtDNA replication intermediates (Fig. S1C-D). We conclude that the Y951N POLγ variant acts in a dominant-negative manner and profoundly impairs mtDNA replication, resulting in both decreased mtDNA copy number and marked replication stalling in cultured human cells.

### POLγ Y951N alteration maintains DNA binding and moderately reduces DNA synthesis activity

To elucidate the mechanism underlying the Y951N-induced mtDNA replication stalling, we purified recombinant WT and Y951N POLγ from insect cells and assessed their DNA binding capabilities (Fig. S2A-C). Electrophoretic mobility shift assay (EMSA) using a TET-labelled DNA substrate showed that the POLγA Y951N variant had a comparable DNA binding affinity to the WT protein (Fig. S2B-C). Furthermore, when paired with the processivity subunit POLγB known to enhance the DNA binding capacity of POLγA, the Y951N mutant performed in a comparable fashion to the WT protein (Fig. S2D).

The position of tyrosine 951 within the active site of the polymerase domain coupled with the observed replication stalling and the unaffected DNA binding properties led us to examine its DNA synthesis activity. We performed primer extension experiments using a radio-labelled 25/70 nt primer/template DNA substrate (Fig. 2A). In contrast to our expectations, the Y951N variant effectively synthesized full-length (70 nt) DNA products, although its DNA synthesis activity was slightly reduced compared to the WT enzyme (Fig. 2B, see lanes 2-4 vs. 8-10). This moderately reduced DNA synthesis by Y951N stands in sharp contrast to the minimal DNA synthesis activity observed upon other active site mutations such as E895G (Fig. 2B, compare lanes 5-7 with 8-10) (30). Next, to mitigate any possible effects of the sequence context during the primer extension reactions, we measured DNA synthesis on a longer DNA template: a circular 3-kb single-stranded pBluescript SK+ with an annealed primer (Fig. 2C). On this longer DNA substrate, the observed small difference in DNA synthesis efficiency of the Y951N mutant was enhanced (Fig. 2C-D, compare lane 1-3 with 4-6); yet no new stall sites were observed, demonstrating that the impairment was not dependent on sequence context.

**Figure 2.**
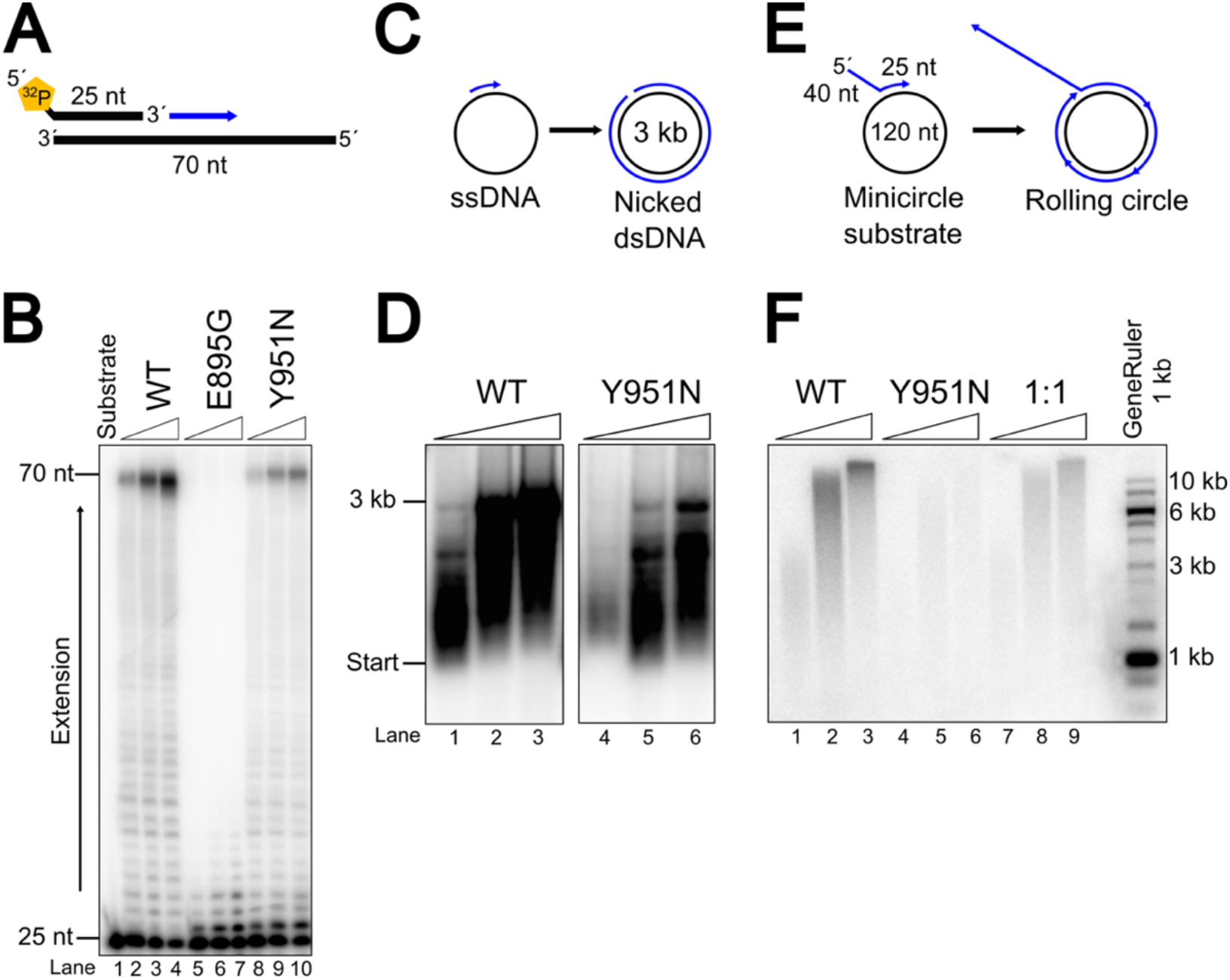
POLγ Y951N displays decreased polymerase activity. **(A)**Schematic overview of the DNA substrate used in (B), a 70 nt template primed with a 5’-radiolabeled 25 nt primer. **(B)**Primer extension with WT POLγ, E895G and Y951N on the primer/DNA (25/70 nt) substrate (shown in A). 12.5 nM POLγ A and 18.75 nM POLγ B_2_ were co-incubated with 5 nM DNA substrate (shown in A) in the presence of 10 μM dNTPs for 20, 40 and 60 seconds. **(C)**Illustration of DNA synthesis on primed 3 kb single stranded DNA pBluescript SK+. **(D)**Primer extension by WT POLγ and Y951N on long DNA substrate (Shown in C). A mixture of 12.5/18.75 nM WT POLγ AB_2_ or Y951N and 750 nM mtSSB was incubated with 5 nM substrate for 10, 30 and 60 min in the presence of 10 μM dNTPs. Radiolabeled α^32^P-dCTP was included to detect the reaction products. **(E)** Schematic overview of the 120 nt minicircle template primed with a 25 nt primer including a 40 nt overhang used in the rolling circle replication assay (shown in F). **(F)** Rolling circle replication by the minimal mtDNA replisome components (12.5/18.57 nM WT POLγ AB_2_ or Y951N, 250 nM mtSSB and 12.5 nM TWINKLE) on a 5 nM minicircle substrate shown in (E) in the presence of 100 µM dNTPs and radiolabeled α^32^P-dCTP. At 1:1 ratio, WT POLγ A and Y951N were added in equal amount to the reaction (6.25 nM each, 12.5 nM in total for POLγ A). Reaction times were 1, 30 and 60 min.

So far, an excess of protein over DNA was used in the assays. Such conditions allow for multiple DNA polymerase rebinding events to occur and could thus potentially mask a reduced processivity for the Y951N variant. To exclude this possibility, we performed DNA synthesis in the presence of heparin which traps all free POLγ molecules, thus preventing any rebinding events (Fig. S3A-B). Even under these single-hit conditions, the Y951N variant was able to extend the DNA substrate by the full 15 nucleotides to the end of the template, but compared to WT POLγ, the distribution of the Y951N products was slightly shifted towards shorter fragments (Fig. S3B, compare lanes 8-10 with 11-13). This experiment shows that the Y951N mutant has a slightly reduced processivity during DNA synthesis.

To achieve maximal processivity when replicating mtDNA, POLγ requires the activity of the mitochondrial DNA helicase TWINKLE and mitochondrial single-stranded DNA-binding protein (mtSSB). To examine the ability of the Y951N mutant to synthesize DNA in the presence of TWINKLE and mtSSB, we used a primed single-stranded mini-circle substrate with a 40-nt 5’-overhang to allow TWINKLE loading. Once initiated, leading strand DNA synthesis by WT POLγ, coupled with continuous unwinding of the DNA template, can in theory continue indefinitely (so-called rolling circle replication; Fig. 2E, lanes 1-3). Compared to WT POLγ, the Y951N variant was substantially impaired in its activity and did not support efficient rolling circle replication (Fig. 2F, compare lanes 1-3 with 4-6). The finding that the DNA synthesis defect of Y951N was exacerbated in the presence of TWINKLE, as opposed to in its absence (when compared to WT POLγ), shows that stimulation by TWINKLE could not rescue the decreased processivity of POLγ Y951N (compare Fig. 2F with 2D). When we combined Y951N and WT POLγ at a 1:1 ratio to simulate the *in vivo* heterozygous state, the polymerase activity of the WT POLγ was only slightly impacted by the presence of the Y951N mutant (Fig. 2F, compare lanes 1-3 with 7-9). We conclude that although the Y951N alteration impairs DNA polymerase activity over longer distances, it does not strongly inhibit the WT POLγ protein, a deviation from the inhibition observed with other dominant active site POLγ mutants such as Y955C (31). Our findings show that the Y951N mutant experiences increasing difficulty with longer DNA sequences, indicating polymerase dysfunction. Moreover, the stimulation by TWINKLE is not sufficient to mitigate the defects in Y951N processivity.

### The Y951N POLγ mutation has an altered balance between DNA synthesis and proofreading

The Y951 residue has been suggested to play a critical role in binding and incorporation of the incoming nucleotide (3, 28, 29). To directly assess the dNTP binding capability of Y951N, we performed filter binding assays in the presence of a DNA primer/template and the complementary incoming nucleotide for the succeeding primer position, [α-^32^P]-dCTP. We used Ca^2+^ as the metal cofactor since it supports the binding of the incoming dNTP to POLγ, yet does not promote primer extension or exonuclease activity. As expected, the Y951N mutant displayed a 2.4-fold reduction in dNTP binding affinity compared to WT POLγ (Fig. 3A), which might explain the reduced DNA synthesis activity. If decreased dNTP binding was the major cause of reduced DNA synthesis, the polymerase activity of Y951N should be restored when increasing the dNTP levels in primer extension reactions. However, in experiments conducted at high dNTP concentrations (up to 1 mM), where dNTP binding is not limiting for DNA synthesis, the Y951N mutant was still unable to produce the same quantities of full-length DNA products as the WT POLγ (Fig. 3B-C, compare lanes 9-11 with 19-21). This result suggests that the decreased affinity for dNTPs might not be the primary cause of the Y951N having lower polymerase activity over longer distances. Even more surprisingly, at very low dNTP concentrations (10 and 50 nM), Y951N was able to produce full-length (40 nt) DNA products while the WT POLγ failed to do so (Fig. 3C, see lanes 3-4 vs. 14-15). These contrasting findings — that Y951N builds more full-length products at low dNTP concentrations but shows reduced DNA synthesis under other conditions — suggests that Y951N may have an altered balance between the polymerase and exonuclease activities. In support of this hypothesis, the Y951N variant demonstrated a decreased capacity to degrade the DNA substrate in the absence of dNTPs (Fig. 3C, see lane 2 vs. 12). This suggests that the Y951N mutation unexpectedly influences exonuclease activity despite being located in the finger domain of the polymerase active site, 32 Å away from the exonuclease active site.

**Figure 3.**
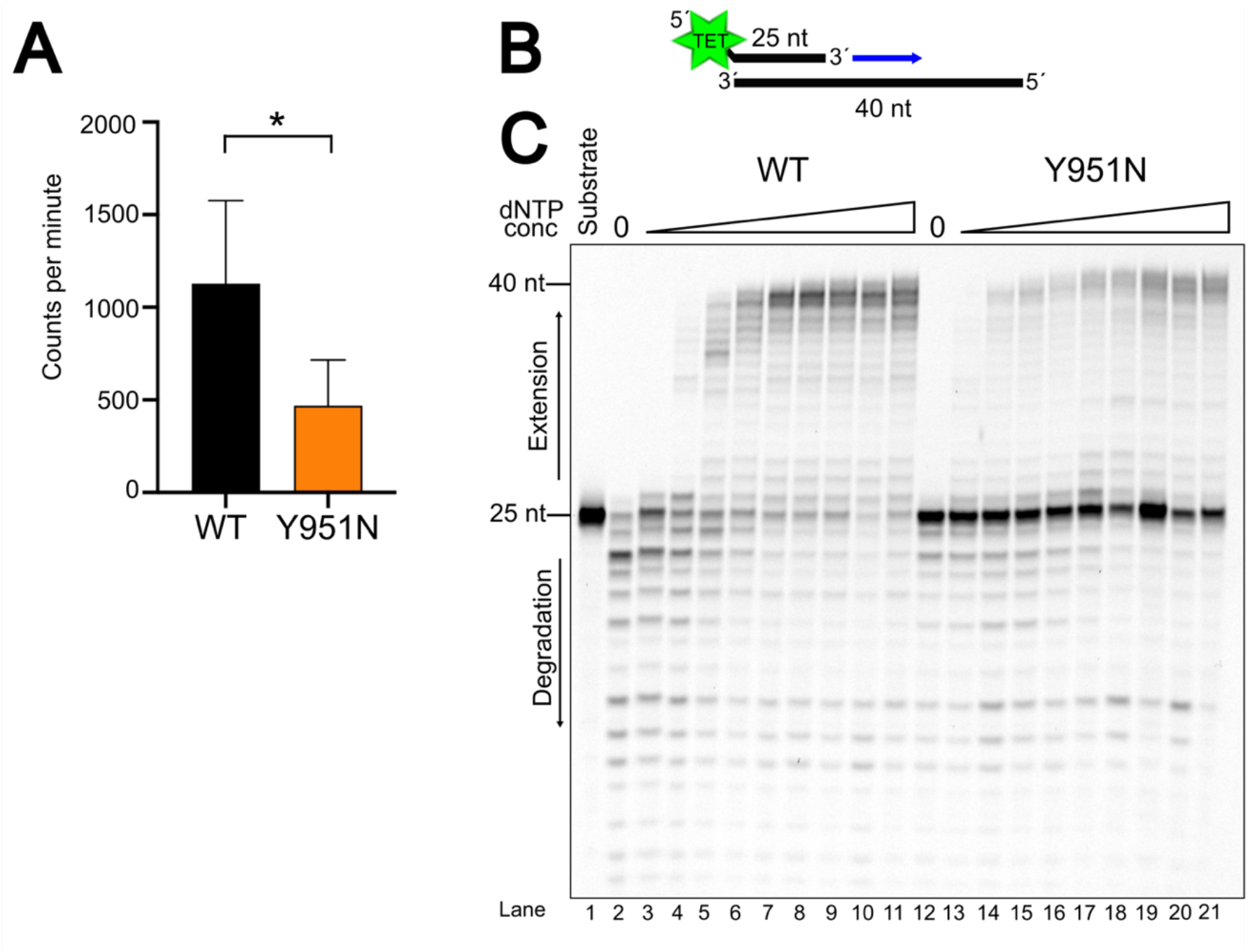
The effect of dNTP concentration on POLγ Y951N DNA synthesis. **(A)**A comparative analysis of dNTP affinity between WT POLγ and Y951N using a filter binding assay. 60 nM POLγ A and 90 nM POLγ B were incubated with 300 nM DNA substrate and 1 μM dCTP including α-^32^P dCTP in the presence of Ca^2+^ for 5 min. After reaction termination, all protein-bound dCTP were caught in a 0.2 μM filter and counts per minute (cpm) were measured using Beckman Coulter LS 6500 Liquid Scintillation Counter. The graph shows the average cpm for WT (1128 cpm) and Y951N (468 cpm) indicating a 2.4-fold decrease in dNTP affinity for Y951N. Error bars represent the standard deviation from six individual replicates. An unpaired t-test with Welch’s correction highlighted a significant difference between WT and Y951N (p = 0.0139). **(B)**Schematic overview of the 5’-TET labeled 25/40 nt primer/template used for the dNTP concentration curve in (C). **(C)**12.5/18.75 nM WT POLγ AB_2_ or Y951N were incubated on 5 nM 25/40 nt substrate with increasing dNTP concentrations (0 nM, 1 nM, 10 nM, 50 nM, 100 nM, 1 µM, 10 µM, 50 µM, 100 µM and 1 mM) and reactions were stopped after 10 min.

### The Y951N is defective in intramolecular polymerase-to-exonuclease switching

To investigate the mechanism by which the Y951N mutation affects DNA degradation, we conducted exonuclease assays on different substrates (Fig. 4A). In contrast to its WT POLγ counterpart, the Y951N variant was hardly able to degrade the partially double-stranded DNA primer/template substrate (Fig. 4B, compare lanes 2-5 with 6-9). Next, we tested exonuclease efficiency of the Y951N mutant using single-stranded DNA (ssDNA) as substrate. Remarkably, the Y951N variant degraded ssDNA as efficiently as the WT POLγ (Fig. 4C). This discrepancy in DNA degradation behavior between the two DNA substrates may be attributed to distinct substrate-specific binding modes. On ssDNA, the DNA polymerase directly enters exonuclease mode upon binding, whereas on partially double-stranded DNA, the protein initially binds in polymerase mode and must first transition to exonuclease mode to initiate DNA degradation.

**Figure 4.**
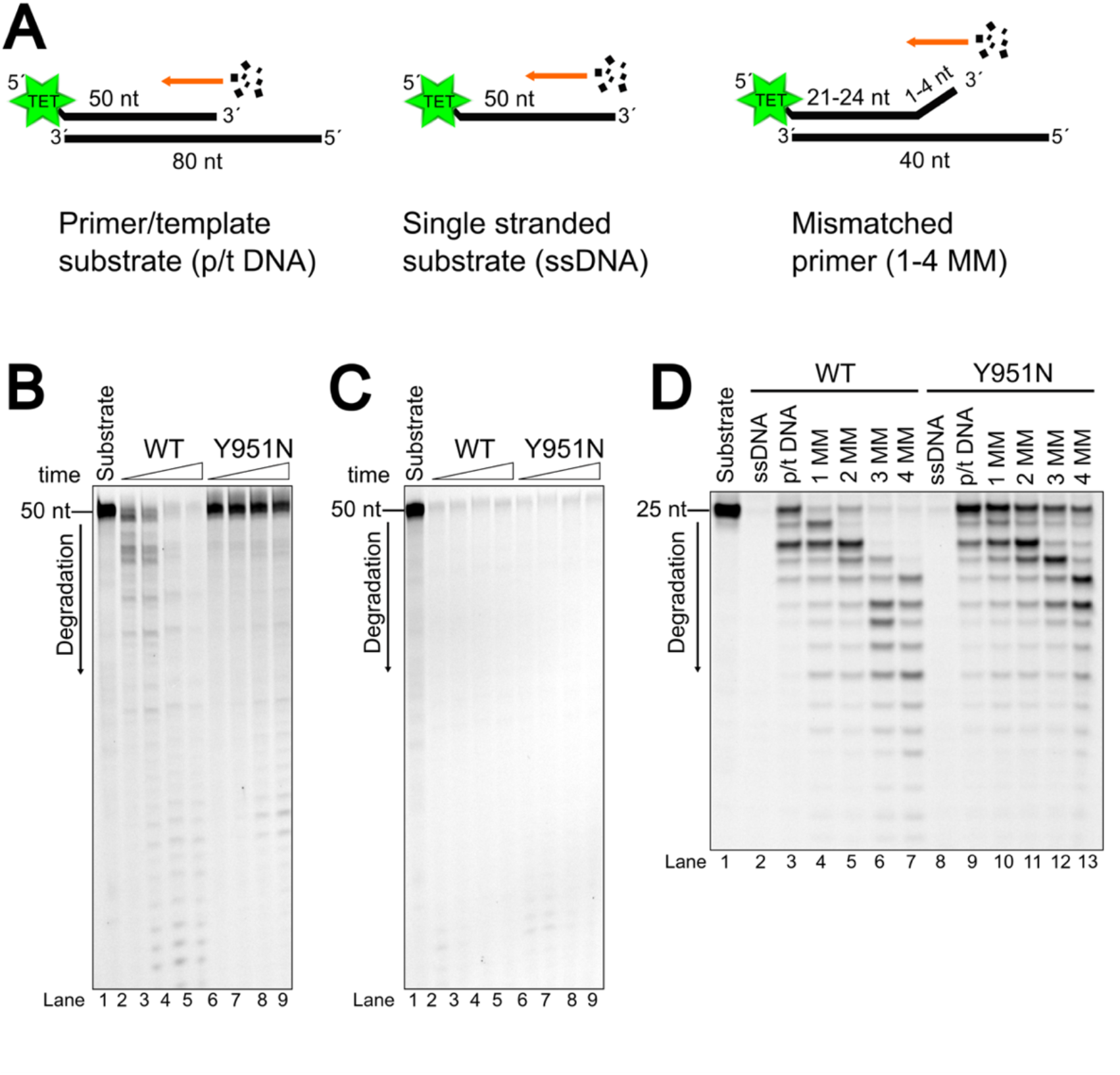
The polymerase-to-exonuclease switch is defective in POLγ Y951N. **(A)**Illustration of the 5’-TET labeled substrates used in exonuclease activity assays in Fig. 4B-D. Left; a partially double stranded 50/80 nt primer/template substrate with a perfectly matched primer (p/t DNA). Middle: a completely single stranded 50 nt substrate (ssDNA). Right: a partially mismatched primer/template with one to four mismatches at the 3’-primer end (1-4 MM). **(B)**Analysis of exonuclease activity on 5 nM partially dsDNA primer/template (left substrate in A) with 25/37.5 nM WT POLγ AB_2_ or Y951N during 5, 10, 30 or 60 min. POLγ Y951N exhibits diminished exonuclease activity compared to WT POLγ on this substrate. **(C)**Assessment of exonuclease activity on a ssDNA substrate (middle substrate in A) by WT POLγ or Y951N during the same condition as in (A). The exonuclease domain of Y951N shows a wild type-like activity on ssDNA substrate. **(D)**Comparative analysis of 30 min exonuclease activities of 25/37.5 nM WT POLγ and Y951N on 5 nM substrates with varying mismatch lengths (right substrate in A); this includes 25 nt ssDNA and 25/40 nt p/t dsDNA as controls and 25/40 nt mismatched dsDNA substrates with one to four mismatches (1-4 MM). Both WT and Y951N POLγ variants are more efficient in exonuclease activity using mismatched substrates compared to complementary primer-template substrate.

Such transitions, typically achieved through an intramolecular switch, are well-documented in various DNA polymerases (14-20). Thus, the degradation of a primer/template DNA substrate is not strictly dependent on the exonuclease activity, but also on the intramolecular switch between the polymerase and exonuclease sites. The observation that Y951N efficiently degrades ssDNA but fails with partially dsDNA (Fig. 4B-C) suggests that Y951N is not exonuclease-deficient, but rather defective in transitioning from polymerase into exonuclease mode. To confirm this hypothesis, we executed exonuclease assays using mismatched primer substrates which can stimulate the DNA polymerase to engage in exonuclease activity more effectively than a perfectly matched primer substrate. As shown in Fig. 4D, increasing the mismatch allowed both WT POLγ and Y951N to more efficiently degrade the primer. Collectively, this data suggests that the Y951N mutation does not impede exonuclease activity per se; instead, it impacts the switching from polymerase to exonuclease mode.

### The Y951N mutation reduces intramolecular transfer of the 3’terminus during exonuclease-to-polymerase transition

During DNA replication, the intramolecular switch not only transitions from polymerase-to-exonuclease mode, but transitions in the reverse direction are also essential after a proofreading event. To investigate if the Y951N POLγ variant is also defective in transitioning from exonuclease-to-polymerase mode, we compared primer extension reactions using mismatched substrates or a complementary primer-template substrate (Fig. 5A). To initiate DNA synthesis on such a mismatched substrate, POLγ should first bind the DNA with its exonuclease site, transferring the 3’-terminus to the polymerase site only after the mismatch is removed. This can be accomplished either by an intermolecular mechanism, where excess polymerase over DNA allows for repeated binding and release of the DNA (Fig. 5B), or by an intramolecular mechanism, demonstrated when polymerases that dissociate from the DNA substrate are prevented from rebinding using heparin as a trap (Fig. 5C). WT POLγ could efficiently extend matched and mismatched primers (Fig. 5B, lanes 2-6). As anticipated, the proofreading-deficient POLγ (EXO^-^) was strongly inhibited on a mismatched primer, reinforcing the importance of exonuclease activity for efficient DNA synthesis on these substrates (Fig. 5B, lanes 13-16). During multiple binding event, the POLγ Y951N variant can extend mismatched substrates to full-length, but it exhibits reduced usage of these substrates compared to the WT POLγ (Fig 5B lanes 7-11).

**Figure 5.**
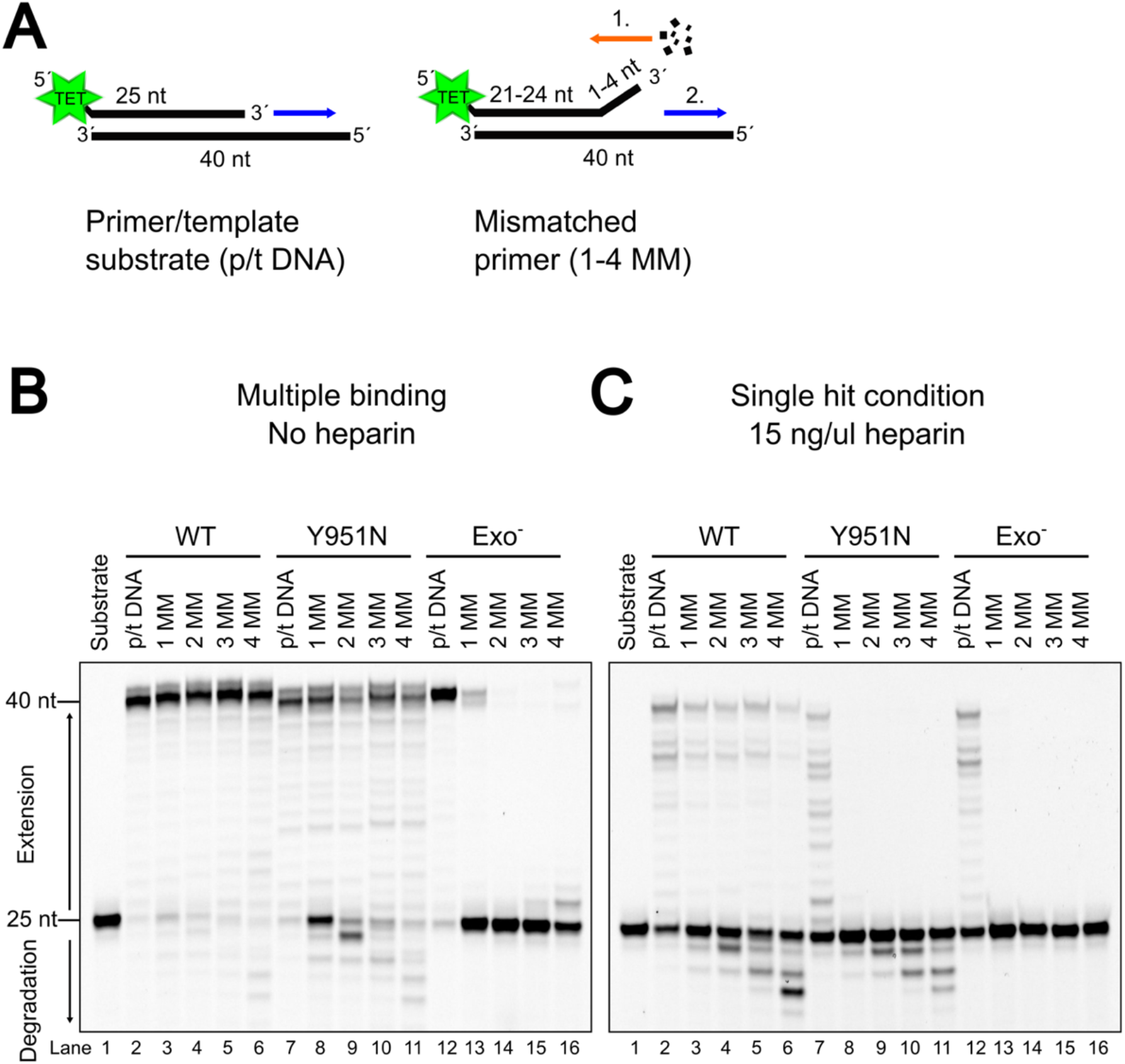
The Y951N mutation leads to an inefficient intramolecular exonuclease-to-polymerase switch in POLγ. **(A)** A comparison of 5’-TET labeled matched and mismatched 25/40-nt primer/template used in primer extension reactions in (B) and (C). The mismatched substrate contained up to four mismatches at the primer’s 3’-terminus. On this mismatched substrate, POLγ is first required to excise the mismatch through its exonuclease activity (1. orange arrow) before extending the primer (2. blue arrow). **(B)** Primer extension reactions comparing matched (p/t DNA) with mismatched substrate (1-4 MM) using either WT, Y951N or exonuclease deficient (Exo^-^) POLγ variants. These reactions were conducted with 100 µM dNTPs, 5 nM substrate and 60/97.5 nM POLγ AB_2_ over a 5-minute duration, under conditions permitting multiple polymerase binding events (no heparin present). **(C)** Primer extension as in (B) but performed in the presence of 15 ng/µl heparin to trap free polymerases, thereby generating “single hit” condition.

In the experiment shown in Fig. 5B, an excess of polymerase over DNA substrate was used, which can favor the usage of an intermolecular switch where the polymerase first binds in EXO mode, dissociates from the DNA, and subsequently rebinds in POL mode. To rigorously examine only the effect of the intramolecular switch from exonuclease-to-polymerase mode, it is essential to prevent these multiple rebinding events. Consequently, we replicated the experiments with heparin present, ensuring only a single binding event per polymerase molecule. The WT POLγ protein, which is proficient at intramolecular switching, managed to remove the mismatches using the exonuclease activity and switch back to polymerase mode to extend the primer on the mismatched substrates with heparin added to the reaction (Fig. 5C, lanes 3-6). As expected, the EXO^-^ POLγ could not extend any of the mismatched DNA substrates. The Y951N mutant also failed to extend these DNA substrates; however, there was notable degradation of the mismatched portion of the primer (Fig. 5C, lanes 8-11). This underscores that while POLγ-Y951N can bind and excise the mismatched segment (step 1, Fig. 5A right panel), it struggles with the intramolecular transition from EXO to POL mode which is needed to extend the primer after mismatch removal (step 2, Fig. 5A right panel).

### The Y951N POLγ alteration leads to decreased intramolecular switching and increased replication stalling during misincorporation events

The experiments using substrates with pre-existing mismatches in the primer (Fig. 4-5) compellingly demonstrate that the Y951N variant has a compromised ability to transition between the EXO and POL sites. However, during DNA replication, DNA polymerases dynamically switch between binding in the POL and EXO sites such that a misincorporation shifts the balance towards the exonuclease site, but the balance tips back in favor of DNA synthesis when the incorrect nucleotides have been removed. To mimic this dynamic POL/EXO switching we performed misincorporation assays in which the DNA polymerase is required to actively transition the 3’-primer terminus between the EXO and the POL site after a running start. For the misincorporation reaction, 1 mM of dATP, dCTP and dTTP were used while dGTP was omitted, forcing a misincorporation opposite a dCMP in the template (position m, Fig. 6C). Once a nucleotide is misincorporated at position m, the polymerase must transition to the exonuclease site to excise the mismatch and thereafter revert to the polymerase site for continued synthesis (Fig. 6A-B). The absence of dGTP will trigger subsequent misincorporation events which leads to additional POL-EXO-POL switching events. Occasionally, a misincorporation is tolerated by the polymerase, leading to extension of the mismatch and continuous DNA synthesis until the polymerase encounters the next dCMP in the template (Fig. 6B).

**Figure 6.**
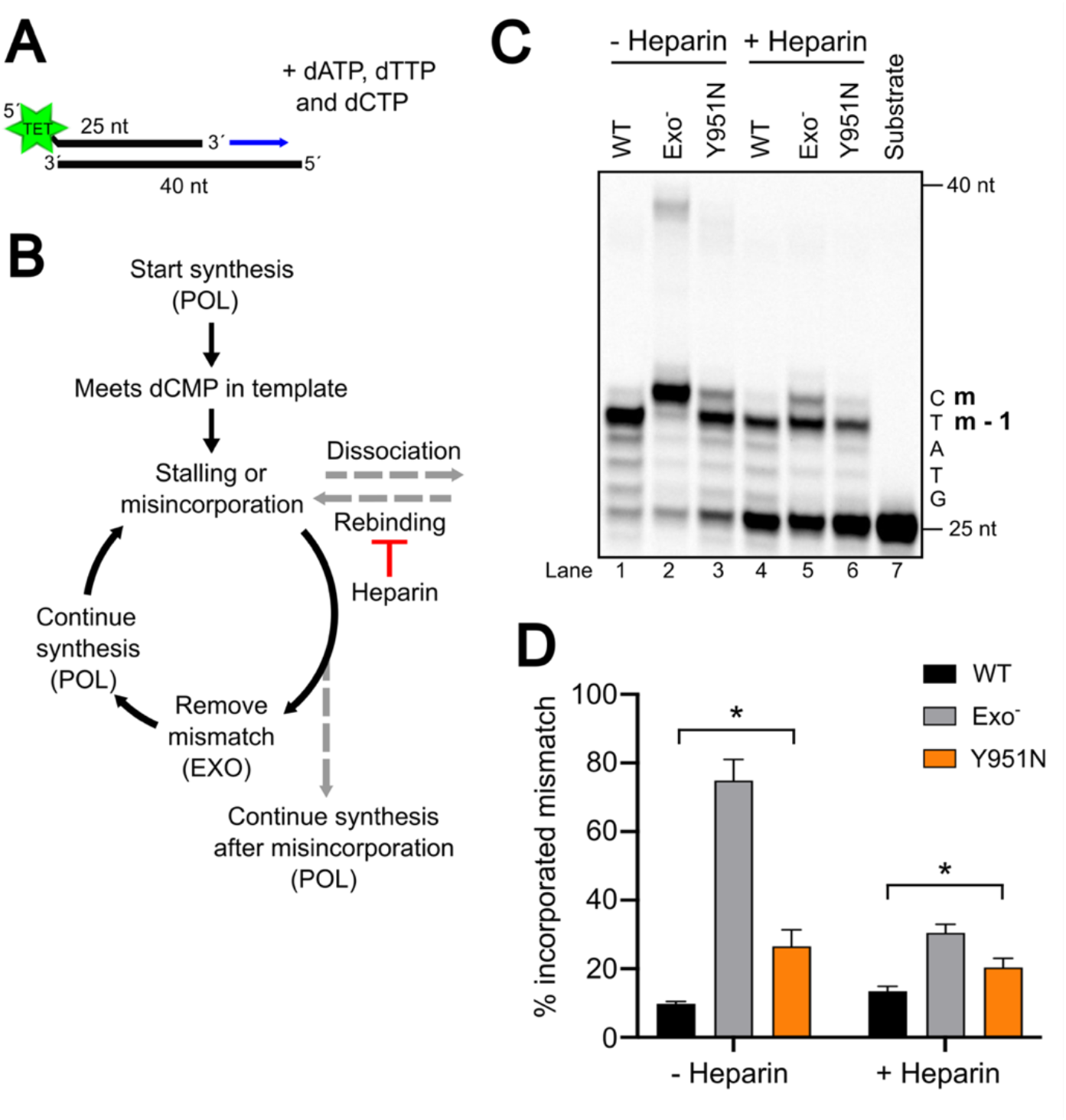
Comparative analysis of mismatch correction by POLγ Y951N than WT during misincorporation. **(A)** Representation of the 5’-TET labeled 25/40 nt primer/template used in the misincorporation assays in (C). **(B)** Schematic illustration of the various kinetic steps within the misincorporation assay, that facilitates the dynamic switching by the DNA polymerase between active modes of the DNA polymerase, denoted by POL (polymerase) and EXO (exonuclease) in capital letters within brackets. Upon dissociation from the substrate, heparin will hinder rebinding of the polymerase to the substrate. **(C)** Misincorporation assays by WT POLγ, Exo^-^ and Y951N variants under conditions that allow multiple binding events (no heparin) or at single hit condition (with 15 ng/µl heparin). 5 nM substrate was incubated with 65/97.5 nM POLγ AB_2_ for 5 min in the presence of 1 mM dATP, dCTP and dTTP. The deliberate exclusion of dGTP induces forced misincorporation at the dCMPs in the template. The **m** indicates the position of the initial mismatch at the first dCMP in the template, with **m-1** marking the position immediately preceding the mismatch. **(D)** Quantification of extended DNA products from (C), band intensities were quantified with the ImageQuantTL software. The proportion of products at position m or beyond, relative to the total extended DNA products (n + 1 and longer), was calculated and expressed as a percentage, indicating the extent of DNA extension past position m-1. Error bars show standard deviation for three individual replicates. Unpaired t-test with Welch’s correction was performed between WT and Y951N in the absence and presence of heparin (p = 0.0245 and p = 0.0259, respectively).

In the absence of dGTP, WT POLγ extended the primer until the misincorporation across dCMP (position “m” in Fig. 6C) where it was able to remove the misincorporated nucleotide, resulting in a dominant DNA product that is shorter by one nucleotide (“m-1”) (Fig. 6C, lane 1 and 4). In the absence of exonuclease activity (POLγ EXO^-^), the polymerase is forced into a misincorporation opposite dCMP and synthesizes longer DNA products than the WT POLγ protein (Fig. 6C, lane 2 and 5). In line with an intramolecular switching defect in the Y951N POLγ variant, the mutant also synthesized longer products than WT POLγ, leading to an increase in DNA products reaching position m and above as compared to WT POLγ (Fig. 6C, compare lane 1 with 3, quantification Fig. 6D). During single-hit conditions where the proteins are restricted to the intramolecular switch, the Y951N misincorporated more often than the WT POLγ variant (20 % vs. 13% respectively; Fig. 6C, lane 4 vs. 6 and Fig. 6D). The results from the misincorporation assay together with the previous *in vitro* data strongly suggest a defect in the dynamic intramolecular switching between the two active sites of the Y951N mutant.

This defect would lead to stronger replication stalling for the Y951N mutant, which is in line with our *in cellulo* data where Y951N-expressing cells demonstrated frequent replication stalling (Fig. 1 and Fig. S1).

## Discussion

In this study, we have explored the impact of the dominant POLγ Y951N patient mutation on mtDNA replication, discovering a critical role of this residue in the maintenance of mitochondrial genome integrity. We demonstrate that the Y951N mutation, identified in a patient with mitochondrial dysfunctions, results in a moderate decrease in DNA polymerase activity. However, the main defect caused by this mutation is the impairment of the enzyme’s ability to efficiently switch between its two distinct modes of action: DNA synthesis (polymerase activity) vs. degradation/proofreading of DNA (exonuclease activity). Overall, this defect leads to replication stalling and mtDNA instability in human cells.

In WT POLγ, the Y951 residue is positioned in the finger domain (in the so-called O-helix), and forms H-bonds with the ?-phosphate and the 3’-OH group of the sugar pucker of the incoming nucleotide. The Y951N substitution leads to reduced interaction between residue 951 and the incoming dNTP due to the smaller size of the asparagine side chain. This likely explains the reduced dNTP affinity and consequently reduced processivity of the polymerase. The reduced affinity for a dNTP, as shown in filter binding assays (Fig. 3A) and replication assays under single-hit conditions (Fig. S3), demonstrates that in the presence of excess level of dNTP the Y951N is less efficient in elongating DNA compared to WT POLγ. The presence of the mitochondrial helicase, TWINKLE, does not rescue the phenotype of Y951N (Fig. 2D, compare lanes 2-3 with 5-6; Fig. 2F).

The 2.4-fold decrease in dNTP binding affinity observed with the Y951N mutation is, however, modest compared to other active site dominant POLγ patient mutations such as E895G and Y955C. These latter mutations in POLγ show substantially higher reduction in dNTP affinity, with E895G displaying a 22-to 92-fold reduction and Y955C a 45-fold reduction in binding affinity for incoming dNTPs (30, 32). This suggests that the Y951N mutation’s detrimental influence, including the accumulation of stalled mtDNA replication intermediates (Fig. S1D) and the rapid loss of mtDNA copy number (Fig. 1D) extends beyond a mere reduction in dNTP affinity, suggesting additional mechanisms in its pathogenicity. Even in experiments where dNTP availability wasn’t limiting, the Y951N mutant produced fewer full-length DNA products than the WT POLγ, suggesting that there are additional mechanistic challenges beyond dNTP binding affinity (Fig. 3C).

For replicative DNA polymerases, there is a delicate balance between the polymerase and exonuclease activities, which can be disrupted if there is a loss of function in one site (10). Typically, a reduction in polymerase activity would lead to an increase in exonuclease activity (33). Despite observing a somewhat reduced polymerase activity in the Y951N POLγ variant (Fig. 2), an increased exonuclease activity is not detected (Fig. 3C and Fig. 4). Instead, our experiments demonstrate that the Y951N mutant is severely defective in switching between the POL and EXO sites, which substantially impacts the overall replication process. This defect in site switching is likely the primary cause of the observed replication stalling and mtDNA instability in human cells.

In conclusion, the characterization of the Y951N mutation in POLγ’s polymerase domain has provided new mechanistic insights in how a mutation in POLγ may have an impact on mtDNA replication fidelity and stability. This mutation disrupts the essential intramolecular switch between POLγ’s exonuclease and polymerase activities, a mechanism vital for the accurate maintenance of the mitochondrial genome. Therefore, the Y951N mutation represents a new mechanism for causing mitochondrial DNA instability, and based on work in other DNA polymerases, it is likely that other pathogenic POLγ mutations may similarly disrupt this critical intramolecular switch.

## Materials and Methods

### Cell culture and mtDNA analysis

Inducible cell lines for the expression of POLγA variants were established using the Flp-In™ T-REx™ 293 host cell-line (ThermoFisher Scientific) as described previously (6). Expression was confirmed using Western blot with anti-flag antibody. For measuring cell growth, cells were seeded in 6-well plate and counted at each indicated time point by staining with Trypan blue dye exclusion (Sigma-Aldrich). Total cellular DNA was isolated using the PureLink Genomic DNA Mini kit for mammalian cells (Thermo Fisher Scientific). MtDNA copy number was determined by multiplex PCR and mtDNA species were visualized using two-dimensional AGE analysis. Further details are provided in SI Materials and Methods.

### Purification of recombinant proteins

The recombinant human mitochondrial replisome components (POLγA and B_2_, helicase TWINKLE and mitochondrial single-stranded binding protein mtSSB) were expressed and purified from Sf9 insect cells as previously described (34, 35). The purified proteins all lacked the N-terminal mitochondrial targeting signal but had an additional C-terminal 6XHis-tag. The three mutant variants of mtDNA POLγA (EXO^-^ D274A, E895G and Y951N) were expressed and purified in similar quantity and purity as WT POLγA.

### *In vitro* replication and exonuclease assays

All *in vitro* experiments were performed in POLγ reaction buffer containing 5-10 nM substrate, 25 mM Tris-HCl pH 7.5, 10 mM MgCl_2_, 100 μg/ml BSA and 1 mM DTT. Nucleotides were included in replication assays at stated concentrations. Unless specified otherwise, 12.5 nM POLγA and

18.75 nM POLγB (dimer) were used. All reactions were performed at 37 °C and terminated at the indicated time by the addition of 0.5 % SDS and 25 mM EDTA. The samples were incubated at 50 °C for 10 min and formamide was added to a final concentration of 47.5 %. The samples were boiled for 5 min and run on a denaturing 10-15 % polyacrylamide gel (7 M urea, 15 % formamide). For radiolabelled samples, the gel was dried and exposed to phosphoimaging in an Amersham Typhoon (Cytiva). For fluorescently labelled samples, the gel, while still between the glass plates, was directly scanned using the Amersham Typhoon imager (Cytiva). Specific details are provided in SI Materials and Methods.

### Filter binding assay

Filter-based dNTP affinity assay was performed with the following reaction condition: 25 mM Bis-Tris propane pH 6.5, 200 µM C_4_H_6_CaO4, 1 mM DTT, 1 µM dCTP (including α-^32^P dCTP, 3000 Ci/mmol, 10 mCi/ml, Hartmann Analytic), 300 nM substrate, 60 nM POLγA and 90 nM POLγB. A 25/40 nt substrate was used with a 3’-ddC on the 25 nt primer to prevent dCTP incorporation.

Reactions were incubated at 37°C for 5 min, loaded on to a 0.2 µm nitrocellulose membrane (Amersham) and washed with 1 ml of 25 mM Bis-Tris propane pH 6.5 and 200 µM C4H_6_CaO4. After washing, the protein-bound α-^32^P dCTP was measured using a Beckman Coulter LS 6500 Liquid Scintillation Counter. The data were analysed using GraphPad Prism software (Graphpad Software Inc., USA) and an unpaired t-test with Welch’s correction was performed to determine significant difference.

## Acknowledgments

The authors wish to thank Sonja Stenmark for technical assistance and Mikael Lindberg at the Protein Expertise Platform (PEP Umeå University) for cloning of pFastbac1 plasmids. Funding for the research conducted in the Wanrooij S. lab was provided by the Knut and Alice Wallenberg Foundation (KAW.2019.0307), and the Swedish Research Council (VR-MH 2018-0278 and VR 2023-02160). The two-dimensional analysis of replication intermediates was funded by the Finnish Academy (projects 332458 and 338227). P.H.W. is supported by the Swedish Research Council (grant number 2019-01874), The Swedish Cancer Society (19 0022 JIA, 22 2381 Pj), The Knut and Alice Wallenberg Foundation (KAW 2021.0053), and The Swedish Society for Medical Research (S17-0023).

