## Supplemental for "The Y951N patient mutation inactivates the intramolecular switch in human mitochondrial DNA POLγ"

**This PDF file includes:**

Supporting Material and Methods
Figures S1 to S3
Tables S1 to S3
SI References

### Supporting Material and Methods

#### Flp-In™ T-REx™ 293 inducible cell lines

The full-length cDNA of POLγA with a C-terminal Flag-tag was cloned into the pcDNA5/FRT/TO vector (Invitrogen) as previously described (1). Site-directed mutagenesis was performed to generate the mutant constructs and all plasmids were validated by Sanger sequencing. The Flp-In T-REx 293 cells were maintained in DMEM high glucose + GlutaMAX (Life Technologies) supplemented with 10 % Fetal Bovine Serum (Sigma-Aldrich), 1 mM sodium pyruvate, 50 µg/mL uridine and Pen/Strep at 37°C in a humidified incubator with 7% $CO_2$ atmosphere. Expression of POLγA variants was induced by adding Doxycycline (Dox) to the growth medium at a final concentration of 100 ng/mL. At this concentration, Dox addition does not lead to mitochondrial toxicity and provides a stable long-term expression of POLγA. For induction periods extending beyond two days, the culture medium was refreshed every second day including the addition of new Dox.

#### Cell growth curves

Flp-In T-REx 293 cells expressing POLγA variants were seeded in triplicate in 6-well plates (5 x $10^3$ cells/well, Cell+, flat base surface, Sarstedt). Cells were treated with 100 ng/mL of doxycycline at time of seeding and cultured at 37°C in a humidified incubator with 7 % $CO_2$ atmosphere. The cultured medium was refreshed every two days. At each indicated time point, cells were imaged with Evos Digital Color Fluorescence microscope (Invitrogen) prior to cell counting. Briefly, cells in triplicate wells were individually harvested by trypsinization and the cells were resuspended in an appropriate volume of medium prior to counting. Cells were counted at 1:2 dilution with trypan blue dye exclusion (Sigma-Aldrich) in an automated cell counter (Countess II FL, Invitrogen). The growth curves were generated in Excel by plotting the total numbers of viable cells as a function of time.

#### Isolation of total cellular DNA and mtDNA copy number analysis

Total cellular DNA was isolated from whole-cell extracts using the PureLink Genomic DNA Mini kit for mammalian cells (Thermo Fisher Scientific). MtDNA copy number was determined by multiplex PCR with 2 ng of total DNA, using Prime Time Gene expression Master Mix (IDT) on a LightCycler96 (Roche). Primers for nuclear and mitochondrial DNA amplification and fluorescent labelled probes (B2M region for nDNA and D-loop region for mtDNA) are listed in Table S1. The PCR program was set as following: initial 10 min at 95 °C followed by 35 cycles of 15 sec at 95 °C, 15 sec at 55 °C and 1 min at 60 °C. Each PCR sample was run in three technical replicates and each condition had at least three biological replicates. Mean Ct values were used to calculate mtDNA copy number relative to non-induced samples using the ΔΔCt method (2).

#### Two-dimensional AGE analysis

Prior to mtDNA extraction, the Flp-In T-REx cells were induced for 24 h with 3 ng/ml doxycycline. MtDNA was isolated from the induced cells and analysed using two-dimensional agarose gel electrophoresis followed by Southern blot as previously described (3, 4). The purified mtDNA was digested with HincII (ThermoScientific) according to manufacturer's recommendation. Southern blotting was performed with a PCR probe detecting the mtDNA non-coding region (nt 35-611) which was labelled with [α-$^{32}$P]dCTP using the Rediprime II random-prime labelling kit (Cytiva).

#### Western blot analysis

Cells were harvested, washed with PBS and lysed in ice-cold lysis buffer (1 % Triton in PBS) for 30 min on ice. The supernatant containing total protein extract was collected after maximum-speed centrifugation for 15 min at 4 °C. The protein samples were determined using the Pierce BCA assay kit (Thermo Scientific). 20 µg of total protein extract was separated on 4-20 % Mini-PROTEAN® pre-cast gels (Bio-Rad) at 150 V and transferred to 0.45 µm nitrocellulose membranes (GE Healthcare Life Sciences) using a Mini-Protean electrophoresis system (Bio-Rad) at 150 mA per gel for 1 h. The membranes were blocked in 5 % non-fat milk for 1 h at RT prior to incubation with primary antibody overnight at 4 °C. The membrane was washed three times in 1X Tris-buffered saline with 0.1 % Tween-20 (TBS-T) and incubated with horseradish peroxidase-conjugated secondary antibodies for 1 h at RT. Following three washes, detection was done using Super Signal™ West Pico Chemiluminescent Substrate (Thermo Scientific) on a ChemiDoc Touch Imaging System (Bio-Rad). Antibody details are listed in Table S2.

**Linear DNA substrates**

Oligonucleotides were purchased from Sigma Aldrich with highest purity grade (PAGE) and are listed in Table S3. Linear substrates for primer extension, misincorporation assays and exonuclease assays were prepared by annealing a TET fluorescent or γ-$^{32}$P 5´-end labelled primer (25 nt or 50 nt) to a linear template (40 nt or 80 nt) in a 1:2 ratio by heating to 95 °C followed by gradual cooling to room temperature.

**Circular DNA substrates**

Single-stranded pBluescript SK+ was purified as described in(5). Primer 682 was annealed to the pBluescript SK+ in a 1:1 ratio.

For the minicircle ssDNA substrates, the ssDNA template was circularized as described (6, 7). Briefly, linear ssDNA (120 nt) was phosphorylated and annealed to a 28 nt oligo with a 14 nt complementary sequence to each side of the linear fragment to facilitate circularization. After ligation with T4-ligase the product was separated on denaturing acrylamide gel, the corresponding band was detected using UV, cut out from the gel, and purified using a Sep-Pak plus C18 column. Afterwards a 65 nt long primer (including a 5´ end 40nt poly T tail) was annealed to the template by heating to 95 °C and slowly cooling to RT.

**Electrophoretic mobility shift assays (EMSA)**

Electrophoretic mobility shift assay was used to assay the DNA-binding of POLγA and $B_2$. A 40 nt template with ddC at the 3´ end was annealed with 5´-TET-25 nt primer at a 1:1.5 primer:template ratio in 100 mM NaCl by incubation at 95 °C for 5 min and cooling down to room temperature. EMSA reaction mixture (15 µL) contained 25 mM Tris-HCl (pH 7.6), 10 mM $MgCl_2$, 1 mM DTT, 100 ng/µL BSA, 300 µM dATP, 10 µM ddCTP, 10 nM of hybridized 5´-TET-primer/template DNA and indicated amounts of purified POLγ variants. In the EMSA reaction with POLγB, the POLγA and POLγB were pre-incubated together for 2 min at RT before DNA substrate was added. After incubation for 10 min at RT, the reaction was stopped by adding 5 µL loading dye (5 mM Tris HCl pH 7.6, 25% glycerol) and kept on ice. DNA-protein complexes were separated on an 6 % polyacrylamide gel in 1XTB buffer at 120V and imaged using an Amersham Typhoon 9400 imager. The percentage of bound DNA was quantified by comparing the intensity of bound DNA to the unbound DNA in a protein-free control reaction.

**Exonuclease activity assays**

Exonuclease assays were performed in the same conditions as the primer extension reactions (described in previous section) but excluding deoxyribonucleotides from the reaction mix.

**Single-hit condition**

For single-hit condition experiments, 5 nM 25/40 nt substrate and 65/97.5 nM POLγA/B were used. POLγ was preincubated on the DNA for 5 min at RT, followed by addition of $MgCl_2$, dNTPs and 15 ng/µl heparin. Heparin titration experiments were performed to determine the optimal heparin concentration that effectively captures all unbound POLγ without hindering the reaction by competing the DNA-bound protein. A control experiment, where POLγ was preincubated with heparin, was consistently performed to exclude any potential rebinding events.

**Misincorporation assay**

The misincorporation assay was performed as previously described in Ganai et al 2015 (see Material and methods "Polymerase to exonuclease switch assay"). Briefly, 130/195 nM POLγ and 10 nM substrate was preincubated in 5 µl buffer A (25 mM Tris-HCl pH 7.5, 100 µg/ml BSA and 1 mM DTT) for 5 min at room temperature. 5 µl buffer B (25 mM Tris-HCl pH 7.5, 100 µg/ml BSA, 1 mM DTT, 20 mM $MgCl_2$, 2 mM dATP, 2 mM dCTP and 2 mM dTTP) or 5 µl buffer C (buffer B with

30 ng/ml heparin) is added to the buffer A. Reactions were incubated for 5 min at 37 °C before terminating reactions by adding of 0.5 % SDS and 25 mM EDTA. Band intensities were quantified using the Image Quant 10.2 software (Cytiva). The percentage of incorporated mismatch was calculated by dividing the band intensity of the mismatch (m) and products above m with the intensity of total extension (n + 1 and above). The data were analysed using GraphPad Prism software (Graphpad Software Inc., USA) and an unpaired t-test with Welch's correction was performed to determine significant difference.

**Rolling circle replication**

Rolling circle replication reaction (50 µl) contained 5 nM primed circular DNA substrate in 25 mM Tris-HCl pH 7.6, 10 mM $MgCl_2$, 1 mM DTT, 100 µg/ml BSA, 4 mM ATP, 100 µM dNTPs and 0.5 µl $\alpha^{32}$P-dCTP (3000 Ci/mmol, 10 mCi/ml, Haatmann Analytic). 250 nM mtSSB was preincubated for 5 min on ice. 12.5 nM TWINKLE, 12.5 nM POL$\gamma$A and 18.75 nM POL$\gamma$B were added, and reactions were incubated at 37 °C. Reactions were stopped at indicated time points by transferring 10 µl into 10 µl of 2X stop solution (1% SDS, 50 mM EDTA in formamide). To remove free $\alpha^{32}$P-dCTP, reactions were cleaned using a Illustra G-25 spin column. Samples were run on a 1 % denaturing agarose gel (1 mM EDTA, 30 mM NaOH) at 20 V for 20 h (at 4 °C). The gel was dried and visualized by phosphorimaging on an Amersham Typhoon system (Cytiva).

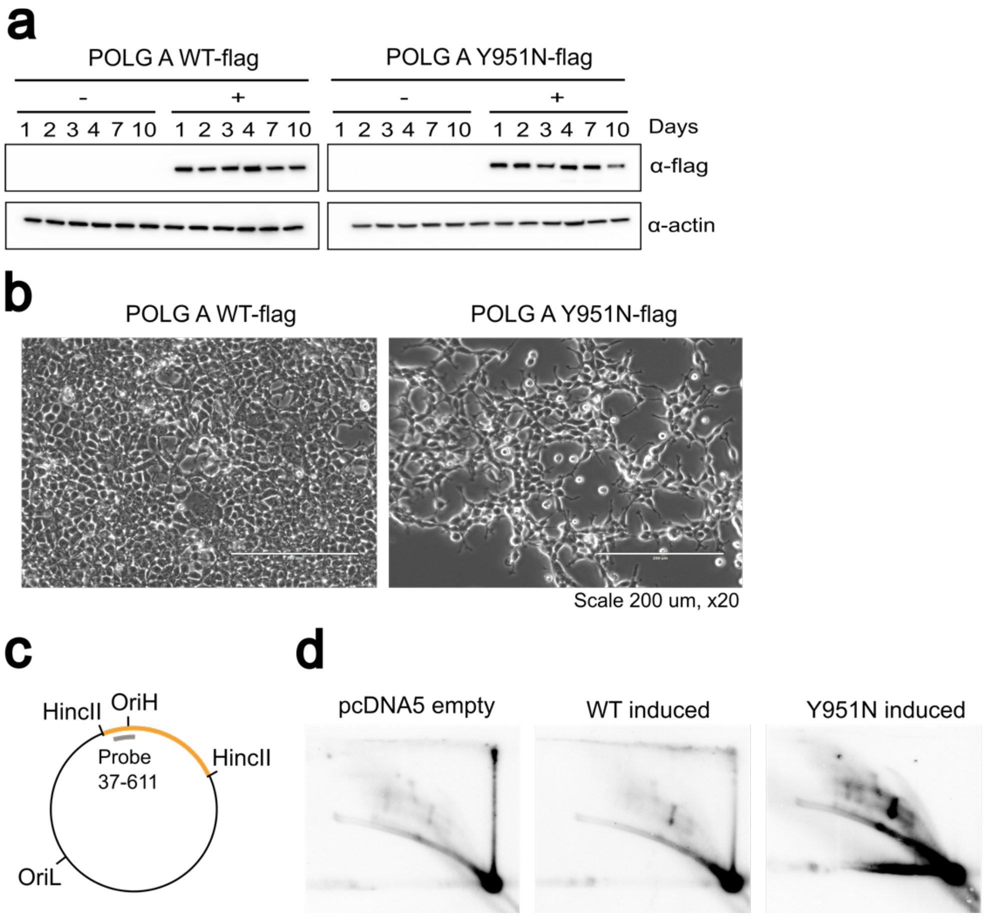

**Fig. S1. Cell growth and mtDNA replication intermediates analysis in the presence of WT or Y951N POLγ variant.**

**(A)** Immunoblot analysis of WT and Y951N POLγ A-flag expressing cells. Anti-flag antibody was used for detection and confirmed the POLγ A-flag expression in the presence of 100 ng/ml doxycycline at the indicated time points. Anti-actin antibody serves as a loading control.

**(B)** Confluency comparison of WT and Y951N POLγ A-flag cells after 10 days 100 ng/ml doxycycline treatment, visualized using the Evos Digital Color Fluorescence microscope (Invitrogen). Cells were imaged prior to cell counting, with a scalebar indicating 200 μm at 20X magnification for size reference.

**(C)** Schematic presentation of human mtDNA with HincII restriction sites (orange) and Southern blot OriH probe (nts 37-611) used for 2D gel analysis indicated in grey.

**(D)** 2D-AGE analysis of mtDNA from parental, WT POLγ A- or Y951N POLγ A-expressing cells probed with OriH probe after 24 h induction of protein expression with 3 ng/ml doxycyline.

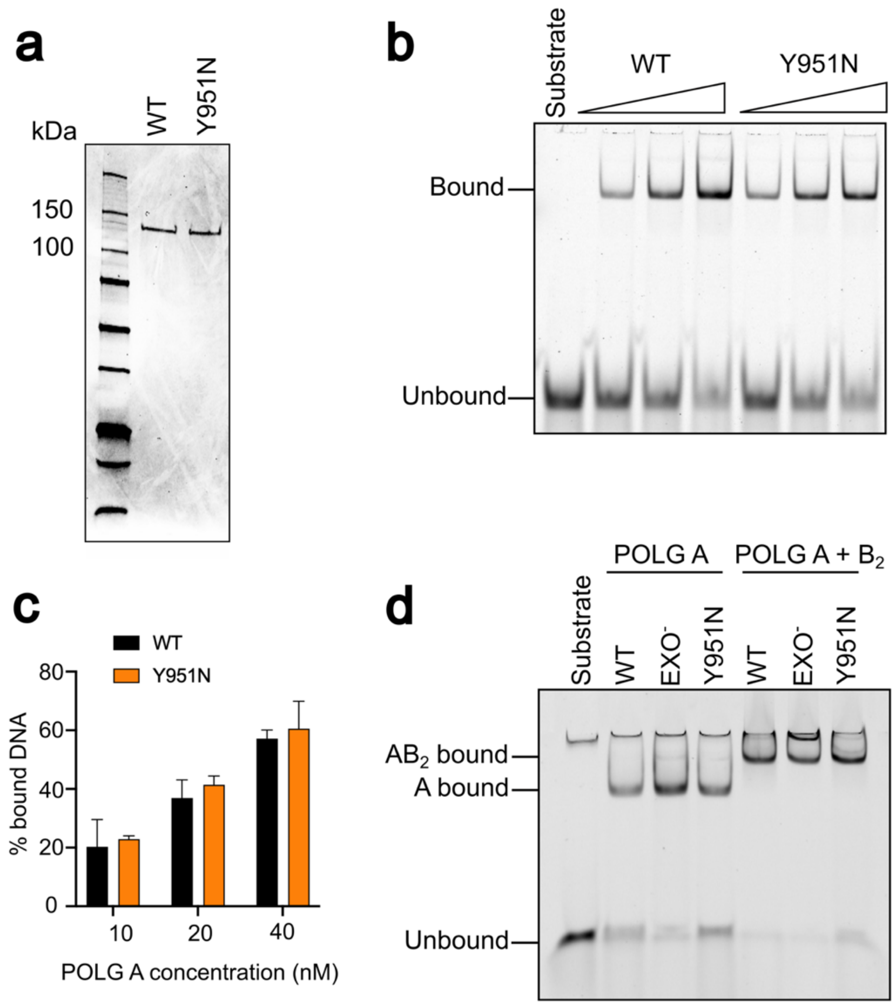

**Fig. S2. DNA binding analysis of recombinant WT and Y951N POLγ variants.**

**(A)** Coomassie brilliant blue stain of purified recombinant WT POLγ A and Y951N POLγ A with 6xHis-tag expressed in Sf9 insect cells. **(B)** Comparative analysis of DNA binding ability between POLγ A WT and Y951N (10, 20 or 40 nM) to a 10 nM TET-labeled 25/40 nt primer-template substrate using EMSA.

**(C)** Quantification of the DNA binding from the EMSA gel in (B). The percentage of bound DNA in correlation with POLγ A concentration. Error bars represent mean -/+ SD of three individual repeats.

**(D)** EMSA of 20 nM POLγ A WT, Y951N or Exo$^-$ together with 30 nM of the processivity subunit POLγ B using a 5 nM 25/40 nt primer-template.

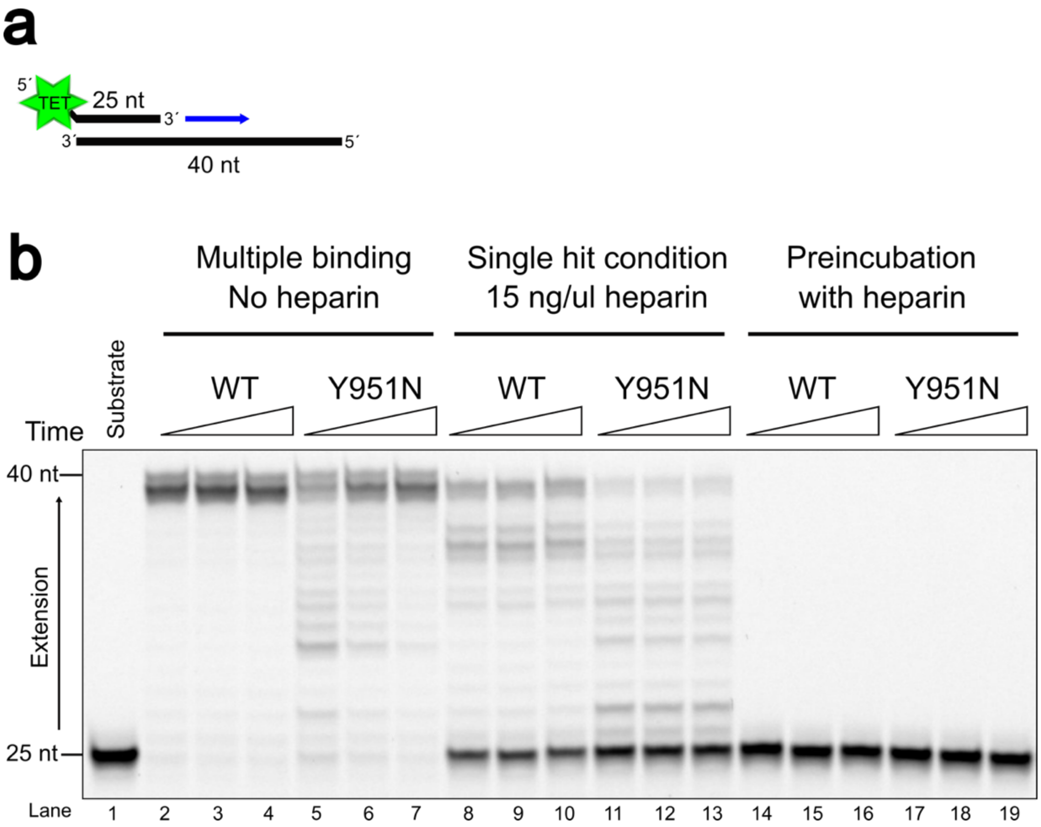

**Fig. S3. Evaluation of WT and Y951N POLγ activity under single hit and multiple binding conditions.**

**(A)** Schematic representation of the 5´-TET labeled 25/40 nt primer/template, used for the optimization of single hit condition with heparin.

**(B)** DNA synthesis by WT POLγ or Y951N in conditions that allow multiple binding events (no heparin), single hit condition (15 ng/µl heparin) or after POLγ and heparin preincubating together. Reactions are stopped at 1, 2 and 5 min.

**Table S1. Primers and probes for multiplex qPCR analysis**

| | | | |
|---|---|---|---|
| Primers | FmtMinArc | mt 16,528 – 16,548 | CTAAATAGCCCACACGTTCCC |
| | RmtMinArc | mt 23 – 42 | AGAGCTCCCGTGAGTGGTTA |
| | Fbeta2M | Chr15 15,798,932 – 15,798,958 | GCTGGGTAGCTCTAAACAATGTATTCA |
| | Rbeta2M | Chr15 15,798,999 – 15,799,026 | CCATGTACTAACAAATGTCTAAAATGGT |
| Probes | PmtMinArc | mt 16,560 – 10 | 6FAM-ATCACGATGGATCACAGGT(NFQ) |
| | Pbeta2M | Chr15 15,798,969 – 15,798,984 | HEX-CAGCAGCCTATTCTGC(NFQ) |

**Table S2. Antibodies used in this study**

| Antibody | Description | Host | Dilution | Manufacturer |
|---|---|---|---|---|
| Anti-flag | Primary monoclonal antibody against flag M2 | Mouse | 1:2000 | Sigma Aldrich F1804 |
| Anti-actin | Primary antibody against beta-actin | Mouse | 1:5000 | Sigma |
| Anti-mouse HRP | Secondary antibody | Goat | 1:20000 | Thermo Scientific |
| Anti-rabbit HRP | Secondary antibody | Goat | 1:20000 | Thermo Scientific |

**Table S3. Oligonucleotides used for *in vitro* assays**

| Description | Sequence 5´- 3´ |
|---|---|
| **For pBluescript SK+** | |
| Primer 682 | TATCGATAAGCTTGATATCGAATTCCT |
| **Rolling circle substrate** | |
| 120 nt ssDNA template | GGCTAGAGGTGGCTAGAATAAATAGGAGGCCT AGGTTGAGGTTGACCAGGGTGTTGTGTATGGT GAGGTGGGTTCATAGTAGAAGAGCGATGGTGA GAGCTAAGGTCGGGGCGGTGATGT |
| Circularization oligo | AGCCACCTCTAGCCACATCACCGCCCCG |
| TET 65 nt primer | TET-TTTTTTTTTTTTTTTTTTTTTTTTTTTTTTTTTTTTTTTTTTTTTTTTTTTT TTTGACCTTAGCTCTCACCATCGCTCTT |
| **For EMSA** | |
| 25 nt ddC primer | TET-GCAGTGAATTTCTGCAGGTCGACT-**ddC** |
| 40 nt ddC template | GCGGGTTGACCTTTGGAGTCGACCTGCAGAAA TTCACTG-**ddC** |
| **Primer extension and exonuclease assays** | |
| 25 nt | ATAGGGGTATGCCTACTTCCAACTC |

| | |
|---|---|
| 70 nt | GAGGGGTATGTGATGGGAGGGCTAGGATATGA GGTGAGTTGAGTGGAGTTGGAAGTAGGCATAC CCCTAT |
| TET 25 nt | TET-GCAGTGAATTTCTGCAGGTCGACTC |
| 40 nt | GCGAGTTGAACTATGGAGTCGACCTGCAGAAA TTCACTGC |
| TET 50 nt | TET-GCTTGTAGACTCCTGCCTAACTGCACAGAGCTG AATTTGCAGTATGGACG |
| 80 nt | GTTATCTAAGCTGCTCTTGGTAGGCATTGACGT CCATACTGCAAATTCAGCTCTGTGCAGTTAGGC AGGAGTCTACAAGC |
| **Mismatched substrate** | |
| 1MM | GCGAGTTGAACTATG**C**AGTCGACCTGCAGAAAT TCACTGC |
| 2MM | GCGAGTTGAACTATG**CC**GTCGACCTGCAGAAA TTCACTGC |
| 3MM | GCGAGTTGAACTATG**CCC**TCGACCTGCAGAAAT TCACTGC |
| 4MM | GCGAGTTGAACTATG**CCCA**CGACCTGCAGAAA TTCACTGC |
